# Mathematical modeling and sensitivity analysis of synNotch-CAR T-cells identify engineering targets for dynamic tunability

**DOI:** 10.64898/2026.03.31.715708

**Authors:** Alexander J. Diefes, Bashir Sbaiti, Maria-Veronica Ciocanel, Cameron M. Kim

## Abstract

Cancer therapeutics are increasingly incorporating engineered receptors due to their ability to detect extracellular ligands and initiate intracellular responses that regulate gene expression. By redesigning these natural signaling systems, synthetic receptors hold great potential for use in novel cell-based therapies. One particularly promising direction is modifying the Notch receptor, a transmembrane protein that naturally mediates ligand-dependent signaling at the cell surface to regulate cell proliferation and differentiation in neurogenesis. Both the intracellular and extracellular domains of Notch can be replaced with alternative domains, creating the family of modified Notch receptors known as synthetic Notch (synNotch). In existing synNotch-activated chimeric antigen receptor (CAR) T-cells, the extracellular domain can be engineered to adjust binding affinity for a specific cancer antigen, enabling precise tuning of therapeutic activity while minimizing off-target effects. To quantify and inform such tuning, we develop differential equations models of synNotch receptor signaling and subsequent gene expression. The mathematical models couple activation dynamics on fast timescales (characteristic of receptor-ligand interactions) and on slow timescales (characteristic of downstream gene expression dynamics). Global Sobol sensitivity analysis of the proposed models highlights parameters that yield the greatest variability in synNotch signal transduction and gene expression, indicating their potential to be engineered for different functions in future cancer therapeutics. For the receptor-ligand interactions in the synNotch model, we find that ligand association and ligand-independent activation are the most sensitive parameters. In the downstream gene expression model, promoter strength and degradation rates of mRNA and gene product are found to be most amenable to engineering.

## 1. Introduction

Chimeric antigen receptor (CAR) T-cell therapy is a promising avenue for treating certain types of cancer, characterized by taking a patient’s own T-cells, engineering them to augment their anti-cancer cell activity with a novel receptor, and reintroducing them back into the patient’s body (1). During the engineering step, these T-cells are modified with a DNA construct that allows them to express a CAR, a synthetic receptor that allows T-cells to recognize and destroy cells expressing a target cancer antigen (2). In the clinic, CAR T-cell therapy has been successful in treating certain types of cancers including relapsed and/or refractory B-cell lymphoma and B-cell acute lymphoblastic leukemia, and these therapies are approved by the U.S. Food and Drug Administration (2–5). However, CAR T-cell therapy has been less effective against solid tumors, as solid tumor antigens are often present on normal tissues at varying levels (2, 6), and solid tumors rarely express one tumor-specific antigen (7). In addition, CAR T-cell therapy is known to trigger cytokine release syndrome, a condition caused by elevated circulating proinflammatory substances, which can be life-threatening if untreated (8, 9). To mitigate these issues, CAR T-cells have been engineered with more sophisticated antigen response systems, often employing other synthetic receptors as part of a genetic circuit to allow the integration of multiple antigen signals. One synthetic receptor used for this purpose is synthetic Notch (synNotch), which is generated by engineering the native Notch protein.

Notch is a cell-surface receptor that plays a role in development and cell differentiation, allowing stem cells to gain specialized function (10, 11). Notch signaling involves direct cell-cell communication, in which a ligand-presenting cell drives the fate of a Notch-presenting cell through physical interactions between Notch receptors and membrane-bound ligands of the ligand-presenting cell (12). Following ligand binding to Notch, the intracellular domain of the protein is cleaved off, and this cleaved fragment serves as a transcription factor for important developmental genes (12). Novel receptors have been designed by swapping functional domains of native Notch to yield the class of receptors known as synNotch (13). The synNotch receptor, like native Notch, contains extracellular, transmembrane, and intracellular domains (13). The extracellular domain binds to cell-surface ligands, and when this domain is modified, the receptor can recognize different ligands and bind at different strengths. Similarly, the intracellular domain can also be swapped so that other transcription factors can be released upon ligand binding (13). Thus, synNotch receptors encapsulate a vast collection of possible combinations of swapped extracellular and intracellular domains, while solely retaining the transmembrane domain that regulates the activation cascade, coupling ligand binding to proteolytic cleavage (13). By leveraging en-gineered domains, input sensing can be tuned extracellularly while the response can also be customized intracellularly, leading to a vast array of possible user-defined sense-and-respond functionalities (13). Since different synNotch receptors with distinct extracellular and intracellular domains do not share signaling intermediates such as an activated kinase or kinase substrates, they are functionally orthogonal, which means that multiple synNotch response programs can be utilized in the same cell without interference (13). This opens the door to creating multifunctional therapeutic cells which utilize Boolean AND/OR-gate logic to require one of or both of two distinct antigens to activate the system (14, 15).

SynNotch has been used in tandem with CAR T-cell therapy in various studies to increase its efficacy, yielding a better tumor killing response, a reduction in off-target effects, or a combination of the two (14–29). The general strategy involves an AND-logic-gated activation pathway that requires two different antigens for the system to activate the killing response: the first antigen activates synNotch, which drives transcription of the CAR, and the second antigen binds to CAR to activate the T-cell (14). This allows for greater specificity, as tumors may exhibit different ensembles of antigens from healthy cells, so this AND-gate logic can be leveraged to target exclusively cancerous cells more reliably than strategies only relying on one antigen (14). SynNotch has also been incorporated into CAR T-cells to generate an OR-gate activation pathway; this is useful when tumors lack both tumor-specific and homogeneously expressed target antigens (29).

When synNotch was first engineered, a significant amount of ligand-independent activation was observed (13, 15). In the realm of cancer therapeutics, this leads to the unintentional activation of immune cells, exacerbating harmful off-target effects. Engineering advances have been made, however, to mitigate this effect. Adding an intracellular hydrophobic sequence found in native Notch was demonstrated to significantly reduce ligand-independent activation, up to 14.6-fold (30). The synNotch receptor has also undergone numerous engineering cycles to achieve different properties such as activation by ligand dimerization (31) and soluble ligand-sensing capabilities (32), which show the continued interest in engineering synNotch. These challenges and advancements highlight the importance of developing predictive quantitative models that will guide synNotch design toward safer and more reliable therapeutic performance.

Due to its vast potential in cellular applications, we seek a thorough mathematical understanding of synNotch receptor dynamics to predict how it can be tuned to achieve desired functions. Mathematical modeling has already been applied to other receptor classes to uncover key mechanistic properties. Ligand-gated ion channels have been studied using mathematical models to generate novel insight in neurology and cardiology (33–41). Enzyme-linked receptors are another class of receptors that have been studied using mathematical modeling for predictions in reaction mechanisms and signal transduction properties (42, 43). The modeling framework we invoke, the cubic ternary complex model (TCM), has been used for ordinary differential equation (ODE) modeling of G protein-coupled receptors, yielding novel insight into agonist/antagonist systems and receptor state interconversions (44–48). Building off of these established modeling frameworks for related systems, we develop a mathematical foundation that can guide engineering for the recently-introduced synNotch receptor.

We are also interested in downstream gene expression, since synNotch is used to activate the CAR gene in synNotch-CAR T-cells. Previously, mathematical models have been successful in predicting the dynamics of gene expression for engineering applications. In the *E. coli* repressilator, an ODE model predicted that a three-gene negative feedback loop could generate oscillations, and a key model parameter constraint was predicted to be fast protein degradation (49). In the *E. coli* bistable toggle switch, another ODE model predicted that two mutually repressing genes could reach steady-states, and it also identified parameters and conditions where cells could flip between steady-states (50). Here, we develop and parameterize a gene expression model for syn-Notch which has the potential to improve our understanding of the synNotch-CAR system.

In Section 2, we develop a synNotch model and a gene expression model, and we describe the Sobol sensitivity analysis method used to assess the global sensitivity of model parameters. We consider the Notch-Delta system as the baseline system, and consider different appropriate parameterizations for different synNotch variants. The gene expression model represents a generic gene product as the base system using a set of parameter values that are in a biologically-relevant range for gene activation and protein translation. In Section 3, we simulate the models and compute the Sobol sensitivity analysis, with a focus on discussing how the results should be considered in an engineering context. In Section 4, we discuss the broader implications of our results and how they can be applied to future engineering efforts.

## 2. Model development and methods

### SynNotch model

We develop a cubic TCM for the molecular synNotch pathways. In the most general model framework, each synNotch receptor can be bound or unbound to ligand, bound or unbound to transcription factor, and be in an activated or inactivated state. These three binary options yield a total of eight possible states. We represent these options by eight corners on a cube, where each spatial dimension of the cube corresponds to a binary option, as shown in Fig. 1a. Traveling upward along an edge of the cube represents binding to a ligand, traveling forward represents releasing a transcription factor, and traveling rightward represents activating the receptor. This framework is consistent with previous applications of cubic ternary complex models (44), although we replace the standard G protein component with the transcription factor, which is Gal4-VP64 for our system (13). The cubic structure allows for consideration of all possible states and transitions of the synNotch system, and the associated differential equations are derived using massaction kinetics.

**Fig. 1.**
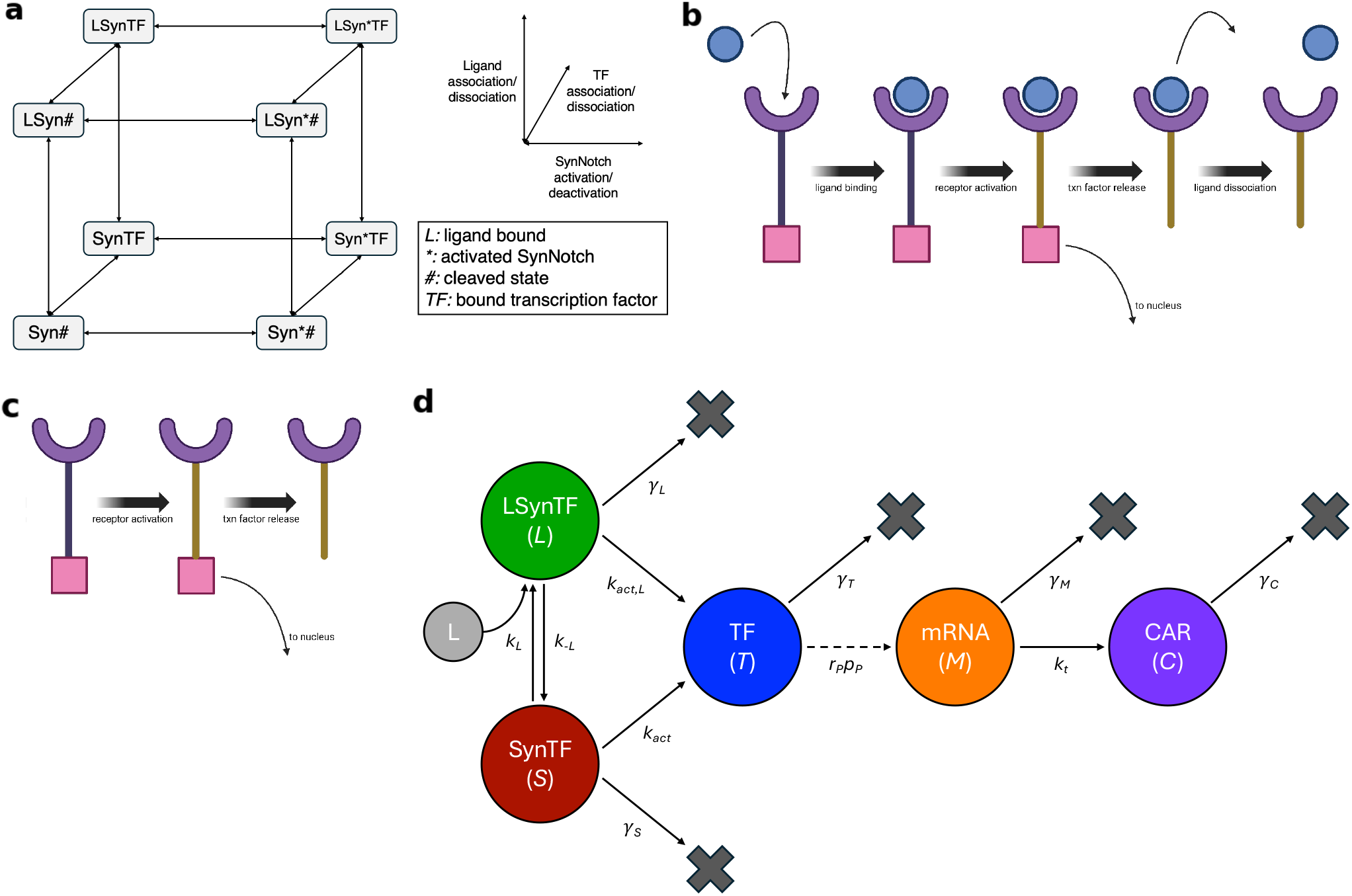
**a**. Cubic ternary complex model schematic for synNotch. Each corner of the cube represents a theoretically-possible state for synNotch. *L* corresponds to ligand bound, ^*^ corresponds to an activated synNotch conformation, # corresponds to a cleaved state with an exposed PEST tag, and *TF* corresponds to transcription factor bound. The PEST tag is exposed once the transcription factor is cleaved and leaves the receptor (13), which signals degradation of the receptor (51). Each orthogonal axis represents a physical property: ligand binding, receptor activation, and transcription factor release. **b**. Primary (ligand-dependent) activation pathway of synNotch. Ligand (blue circle) binds to the extracellular domain of the synNotch receptor (purple curve), which induces activation of the receptor (yellow line). The transcription factor (pink square) is then cleaved and sent to the nucleus. The ligand then dissociates and the receptor eventually degrades. The binding, activation, and dissociation reactions are characterized by the rate constants *k*_*L*_, *k*_*act,L*_, and *k*_−*L*_, respectively; the release of transcription factor is treated as a quasi-steady-state approximation. **c**. Ligand-independent activation pathway of synNotch. The cartoon components are the same as those in the ligand-dependent activation pathway. The receptor activates in the absence of ligand at rate *k*_*act*_. **d**. Schematic of the relevant receptor states and kinetic pathways in the cubic TCM and gene expression model. The variables and parameters for the cubic TCM are given in Table 1, and the variables and parameters for the gene expression model are given in Table 2.

Based on documented biological phenomena, we make four assumptions that allow us to simplify this framework into a minimal model with fewer parameters. First, we assume that once a transcription factor is cleaved, it cannot return to the synNotch receptor as this is a covalent bond (13). This removes the bidirectionality in the backward direction of the schematic in Fig. 1a. Second, we assume that synNotch activation is irreversible, which has been demonstrated for the Notch-Delta canonical pathway, the base system (52). This reduces the leftward direction of the schematic. Third, we assume that activated synNotch with a transcription factor bound is transient, since the transcription factor is released by cleavage upon completion of activation. This removes the *Syn*^*^*TF* and *LSyn*^*^*TF* states. Finally, we assume that the transcription factor cannot release without receptor activation, since it relies on a conformational change (53–55). This reduces the forward pathways from *LSynT F* to *LSyn*# and from *SynTF* to *Syn*#, so the *LSyn*# and *Syn*# states are also removed. Since we do not differentiate between the transcription factor released from the ligand-bound and from the ligand-unbound synNotch, we can combine the *LSyn*^*^# and *Syn*^*^# to a single state, which we refer to as *TF*.

These assumptions simplify the cubic TCM into a three-state system, in which two primary pathways are responsible for releasing the transcription factor: ligand-dependent and ligand-independent activation. The ligand-independent activation pathway is known to be nontrivial, and efforts are being made to diminish it (13, 15, 30). The three states in the reduced system are unbound synNotch (*S*), ligand-bound synNotch (*L*), and released transcription factor (*T*). Table 1 describes the variables and parameters used in the reduced cubic TCM. The canonical ligand-dependent pathway car-toon is shown in Fig. 1b, and the ligand-independent pathway is illustrated in Fig. 1c. The reduced three-state system is shown in Fig. 1d, which also shows its relationship to the gene expression model. The reduced system of differential equations for these three states is given by:

**Table 1.** Variables and Notch-Delta baseline parameters used in solving the cubic TCM.

| Symbol | Description | Units/Value | Reference |
| --- | --- | --- | --- |
| **Variables** |  |  |  |
| $S$ | Unbound synNotch receptor | molecules/cell | – |
| $L$ | Ligand-bound synNotch receptor | molecules/cell | – |
| $T$ | Released Gal4-VP64 transcription factor | molecules/cell | – |
| **Parameters** |  |  |  |
| $k_L$ | Notch-Delta association | $0.167 \mu\text{m}^2\text{s}^{-1}$ | Khait et al., 2016 (56) |
| $k_{-L}$ | Notch-Delta dissociation | $0.034 \text{s}^{-1}$ | Khait et al., 2016 (56) |
| $k_{act,L}$ | Notch-Delta ligand-dependent activation | $0.34 \text{s}^{-1}$ | Khait et al., 2016 (56) |
| $k_{act}$ | Notch-Delta ligand-independent activation | $0.00034 \text{s}^{-1}$ | Estimated |
| $\gamma_S$ | Notch degradation | $0.1 \text{hr}^{-1}$ | Sprinzak et al., 2010 (57) |
| $\gamma_L$ | Notch degradation | $0.1 \text{hr}^{-1}$ | Sprinzak et al., 2010 (57) |
| $\gamma_T$ | Gal4 degradation | $0.265 \text{hr}^{-1}$ | Sprinzak et al., 2010 (57) |

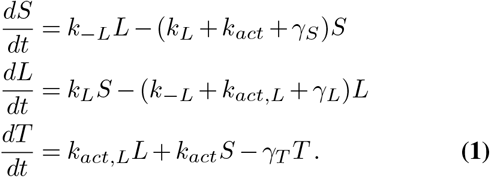

Solving this system allows us to predict the evolution of the relevant states over time.

### Gene expression model

We also develop a model for gene expression, which takes input from the function *T* (*t*), describing the amount of transcription factor inside the cell in Eq. (1). The model we develop here captures gene expression for some generic gene product; for synNotch-CAR T-cells, this product is often the CAR. However, we do not use CAR-specific parameters, as CARs can be engineered to have different properties, like synNotch (58); if we were to explicitly model the product and its activity, we would be more concerned with the baseline values of these parameters. We therefore use a set of parameter values that are in a biologically-relevant range for gene activation and protein translation. We later discuss considering a large range of parameters in our sensitivity analysis so that the exact protein-specific parameter values are not as important.

This model accounts for the two most significant rate-limiting steps involved in protein production: transcription and translation (59). We choose not to explicitly model the trafficking of transcription factors to the nucleus and of mRNA to the endoplasmic reticulum, because these trafficking processes operate on fast timescales, on the order of a few seconds (60, 61). On the other hand, transcription and translation create single protein copies on the order of minutes to hours (59). Thus, we derive time-dependent equations for the number of mRNA molecules *M* (*t*) per cell and translated CARs *C*(*t*) per cell. These variables are illustrated in Fig. 1d.

We use methods from statistical thermodynamics to represent mRNA production by transcription (70). With this approach, the general differential equation that describes the time-dependent amount of mRNA expressed *M* (*t*) takes the form

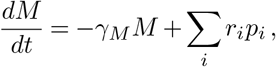

where each transcriptionally active state has a rate of production (transcription) *r*_*i*_ and a probability of occurrence *p*_*i*_. The *γ*_*M*_ *M* term models the degradation of mRNA via first-order kinetics. In our case, we only consider one transcriptionally active state, which is the state where RNA polymerase (RNAP) is bound to the promoter. We denote the transcription rate with bound RNAP by *r*_*P*_ and the probability that RNAP is bound by *p*_*P*_, so this equation becomes

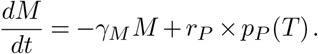

The promoter activity is modulated by the presence of transcriptional activator Gal4-VP64, a fusion protein made by joining the DNA-binding and dimerization domains of Gal4 to four copies of the powerful transcriptional regulator VP16 (15). Gal4-VP64 activates transcription by encouraging the recruitment of RNAP, so *p*_*P*_ is a function of the transcription factor concentration in the cell.

A previously-reported statistical dynamics derivation (69) suggests that we can assume that *p*_*P*_ (*T*) takes on the following functional form:

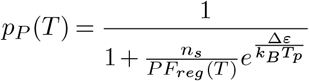

where the regulation factor (*F*_*reg*_) is also a function of *T*. Table 2 describes the variables and parameters used in the gene expression model. The regulation factor *F*_*reg*_ models how the effective number of RNAPs available for binding changes as the transcription factor binds to DNA. Intuitively, the more transcription factor that binds to the DNA, the more RNAPs that are actively recruited. In our case, the exact mechanism by which Gal4-VP64 activates gene expression in a cooperative manner across many Gal4 upstream activating sequence (UAS) sites is unclear. Some theories state that the VP64 domain interacts with different locations on the transcriptional initiation machinery, causing cooperative activation (71). Others explain the cooperativity with chromatin unwinding (72). Due to the lack of a mechanistic understanding of the Gal4-VP64 cooperatively binding to DNA, we use an empirical model to capture this process. We propose a Hill function with the following form:

**Table 2.** Variables and parameters used in solving the gene expression model.

| Symbol | Description | Units/Value | Reference |
| --- | --- | --- | --- |
| **Variables** |  |  |  |
| $T$ | Released Gal4-VP64 transcription factor | molecules/cell | – |
| $M$ | Transcribed mRNA | molecules/cell | – |
| $C$ | Translated CAR | molecules/cell | – |
| **Parameters** |  |  |  |
| $\gamma_M$ | Degradation rate of mRNA transcript | $0.0012/60 \text{s}^{-1}$ | Yang et al., 2003 (62) |
| $\gamma_C$ | Degradation rate of expressed protein | $2.5 \times 10^{-4}/60 \text{s}^{-1}$ | Schwanhäusser et al., 2011 (63) |
| $r_{ext}$ | Extension rate of RNAP II | $3.5/60 \text{kb s}^{-1}$ | Fuchs et al., 2014 (64) |
| $l$ | Expressed protein transcript length | 0.8 kb | Assumed |
| $n_s$ | Number of nonspecific binding sites for RNAP | $10^9$ | Wunderlich et al., 2009 (65) |
| $P$ | Total number of RNAP available for binding per cell | 140000 | Kimura et al., 1999 (66) |
| $F_{reg}$ | Regulation factor (function of $T$) | Varies | – |
| $\Delta\varepsilon$ | Specific and nonspecific binding energy difference of RNAP to DNA | $-3.74 k_B T_p$ | Calculated |
| $k_B$ | Boltzmann constant | $1.38 \times 10^{-23} \text{J K}^{-1}$ | – |
| $T_p$ | Physiological temperature | 310.2 K | – |
| $\alpha$ | Fold activation ($T = 0$ to $T \rightarrow \infty$) | 110 | Calculated |
| $K_{d,T}$ | Dissociation constant estimated based on the $K_d$ of Gal4 dimer binding to one UAS site | 0.1 nM | Luo et al., 2014 (67) |
| $n_T$ | Hill coefficient assumed since Gal4 binds as a dimer to each of 5 UAS sites | 5 | Gadhavi et al., 1996 (68) |
| $k_t$ | Translation rate per mRNA | $2.3/60 \text{copies (mRNA}\cdot\text{s)}^{-1}$ | Schwanhäusser et al., 2011 (63) |
| $r_P$ | mRNA production rate with bound RNAP | $\frac{3.5}{60 \cdot 0.8} \text{mRNA s}^{-1}$ | Calculated |
| $S_0$ | Number of synNotch receptors | 10000 | Assumed |
| $p_P$ | Probability that RNAP is bound (function of $T$) | Varies | Bintu et al., 2005 (69) |

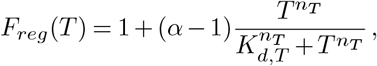

where *α* represents the maximum fold change in the amount of RNAP available when the transcription factor is saturated, *K*_*d,T*_ is the effective dissociation constant (transcription factor concentration such that activation is at 50% of the maximum), and *n*_*T*_ is the Hill coefficient.

Thus, *F*_*reg*_ requires the concentration of transcription factor as an input. The cubic TCM predicts *T* (*t*) (Eq. (1)), and we convert this quantity to nM in the nucleus. We use 125 fL as the average volume of T-cell nuclei, which was previously estimated by taking an average nuclear radius of 3.1 *µ*m, assuming that nuclei are spherical (63). We derive the conversion factor using the assumption that the transcription factor trafficks into the nucleus almost instantaneously relative to the long timescales in which mRNA and protein are expressed. The mRNA and protein concentrations are later converted back to units of molecules/cell for consistency using:

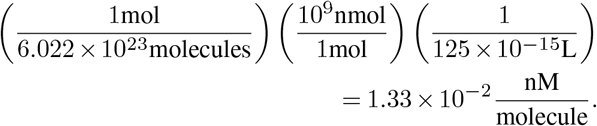

Now, we consider the amount of gene product expressed over time, *C*(*t*). This product is created by translating mRNA, and it also degrades via first-order kinetics. We include the differential equation for *C*(*t*) in the following system, which we henceforth refer to as the gene expression model:

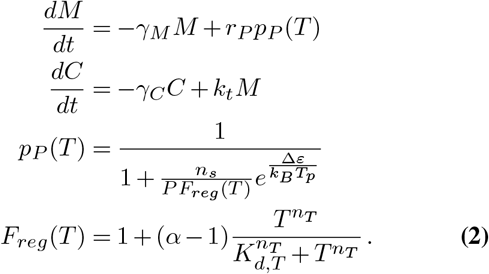

### Sobol sensitivity analysis

The outputs of the synNotch model (Eq. (1)) and gene expression model (Eq. (2)) are the released transcription factor *T* (*t*) and the expressed gene product *C*(*t*), respectively. Modifying parameters in these models could lead to changes in these relevant quantities, and having a quantitative understanding of these relationships would allow engineers to make informed decisions about which components to engineer for improved properties. Our goal is to predict the parameters that are responsible for the largest proportion of the variation in the outputs of each of these models, as these suggest targets for engineers to produce different dynamical responses through protein modifications. We aim to use a global method of sensitivity analysis, since these approaches allow for varying multiple parameters at a time across large domains of the parameter space, which in turn allows to account for individual parameter effects as well as interactions between parameters and how these interactions affect the output (73). Here, we use the Sobol sensitivity analysis method, since it does not require a linear or monotonic relationship between model parameters and output (73, 74). A disadvantage of this method is its high computational cost (73).

Both model outputs we are interested in correspond to protein amounts over time; however, the Sobol sensitivity method requires a scalar measure. First, we specify a time interval for each solution *T* (*t*) and *C*(*t*), denoted as [*a, b*] and [*c, d*] in Eq. (3) and Eq. (4) below, to capture solution behavior. The values of these parameters are discussed and justified in Section 3. For *T* (*t*), we choose *a* and *b* such that the *T* (*t*) solution achieves a steady state, so that this interval corresponds to when most transcription factor has been released. For *C*(*t*), we choose *c* and *d* such that *C*(*t*) has reached a maximum value and *M* (*t*) has reached an approximate steady-state of zero, so there are no mRNA molecules left to be translated into CARs. The choices of these parameters do not have to be exact, since they primarily serve as a consistent measure for comparison when evaluating the model for each Sobol sample of the parameter space. Based on these time intervals, we define the measures using the *L*^2^-norm value of the outputs as follows:

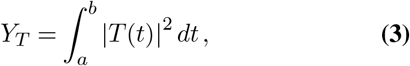

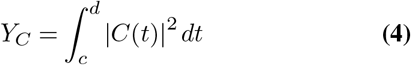

These measures, *Y*_*T*_ and *Y*_*C*_, thus capture characteristics of the dynamics of *T* (*t*) and *C*(*t*) respectively and output scalar values that can be compared under different parameter conditions. We give a brief description of Sobol sensitivity analysis in the Supplementary material. This analysis returns two measures of sensitivity for each parameter: *S*_1_, the first-order sensitivity index, gives a measure of how much the given parameter influences the variance in the output measure, while *S*_*T*_, the total sensitivity index, gives a measure of how much the given parameter and its interactions with other parameters influence the variance in the output measure.

## 3. Model simulations and sensitivity analysis

### SynNotch model and variants

We first simulate the synNotch model in Eq. (1) using Notch-Delta baseline parameters as shown in Table 1. We consider the initial conditions:

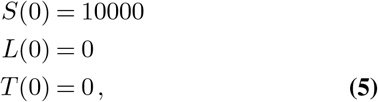

which reflect the scenario where 10000 synNotch receptors are expressed on the surface of a cell and the ligand is first introduced into the system. We also assume that there is a large pool of ligand present so that the association rate is not affected by depletion of available ligand. The cubic TCM simulation yields the predicted protein concentrations in Fig. 2. The number of synNotch molecules in the unbound state drops as the ligand binds, and the transcription factor is subsequently released. The system saturates at approximately 30 seconds of simulation.

**Fig. 2.**
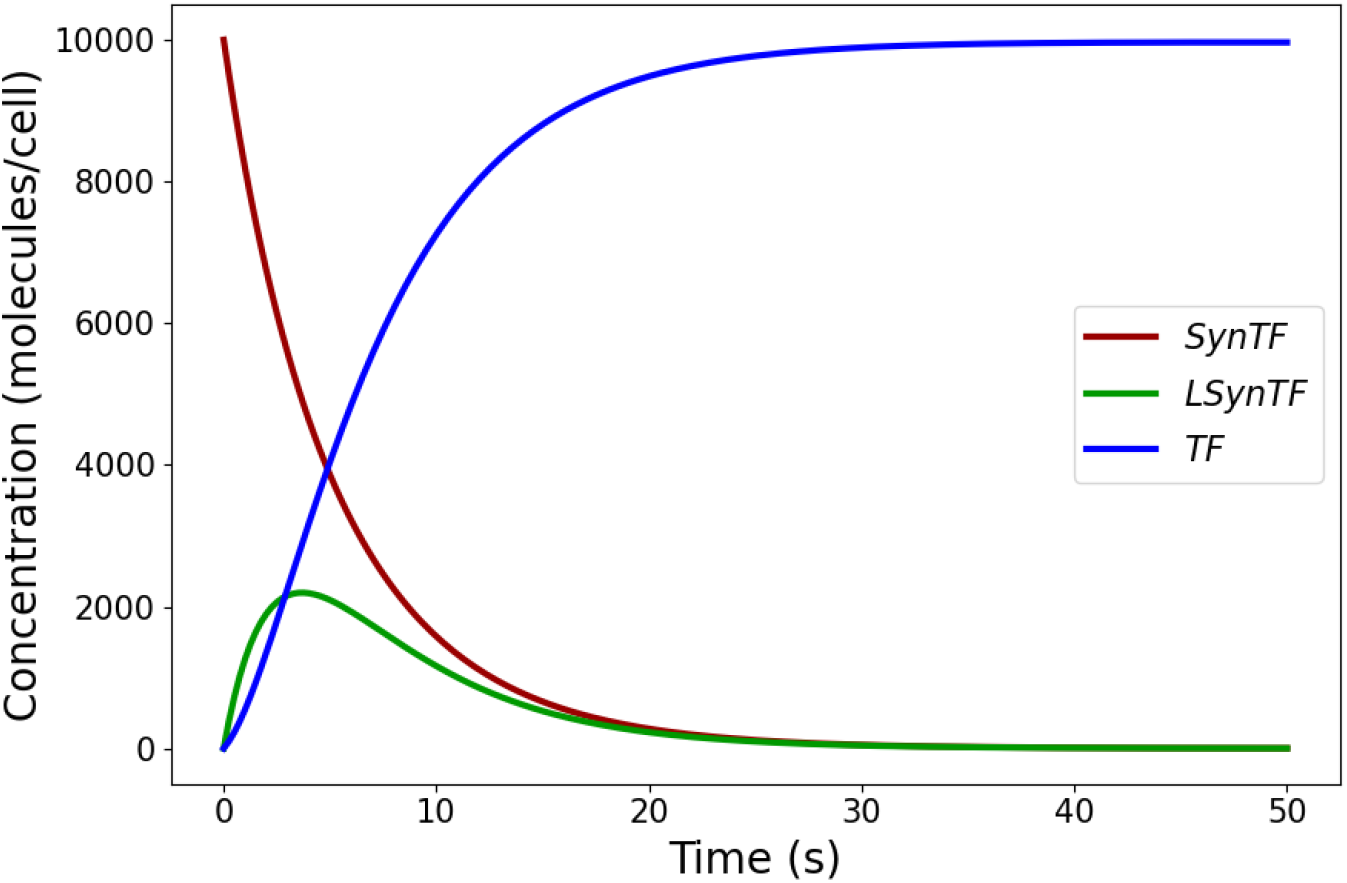
Solution of the cubic TCM with Notch-Delta baseline parameters (Table 1).

We then use the *L*^2^-norm of the transcription factor *T* (*t*) output (from 0 to 50 seconds) as the measure for Sobol sensitivity analysis as in Eq. (3). In the analysis, we choose parameter sets at random on a logarithmic scale in intervals between three orders of magnitude below and three orders of magnitude above their baseline values. We use 2_14_ samples from these intervals for assessing parameter sensitivity. This allows for a broad exploration of the parameter values, thus incorporating predictions for how different kinetic alterations to synNotch could change the transcription factor release dynamics. The results of the sensitivity analysis are shown in Figs. 3a,b, and they reveal that the most sensitive parameters are ligand association (*k*_*L*_) and ligand-dependent activation (*k*_*act,L*_). It should also be noted that the *S*_1_ indices and the *S*_*T*_ indices differ, so higher-order interactions between the parameters contribute to the variation in output. Overall, parameters with higher total Sobol indices could represent targets for engineering, since changing their corresponding components of the synNotch receptor may yield the most significant changes in released transcription factor dynamics.

**Fig. 3.**
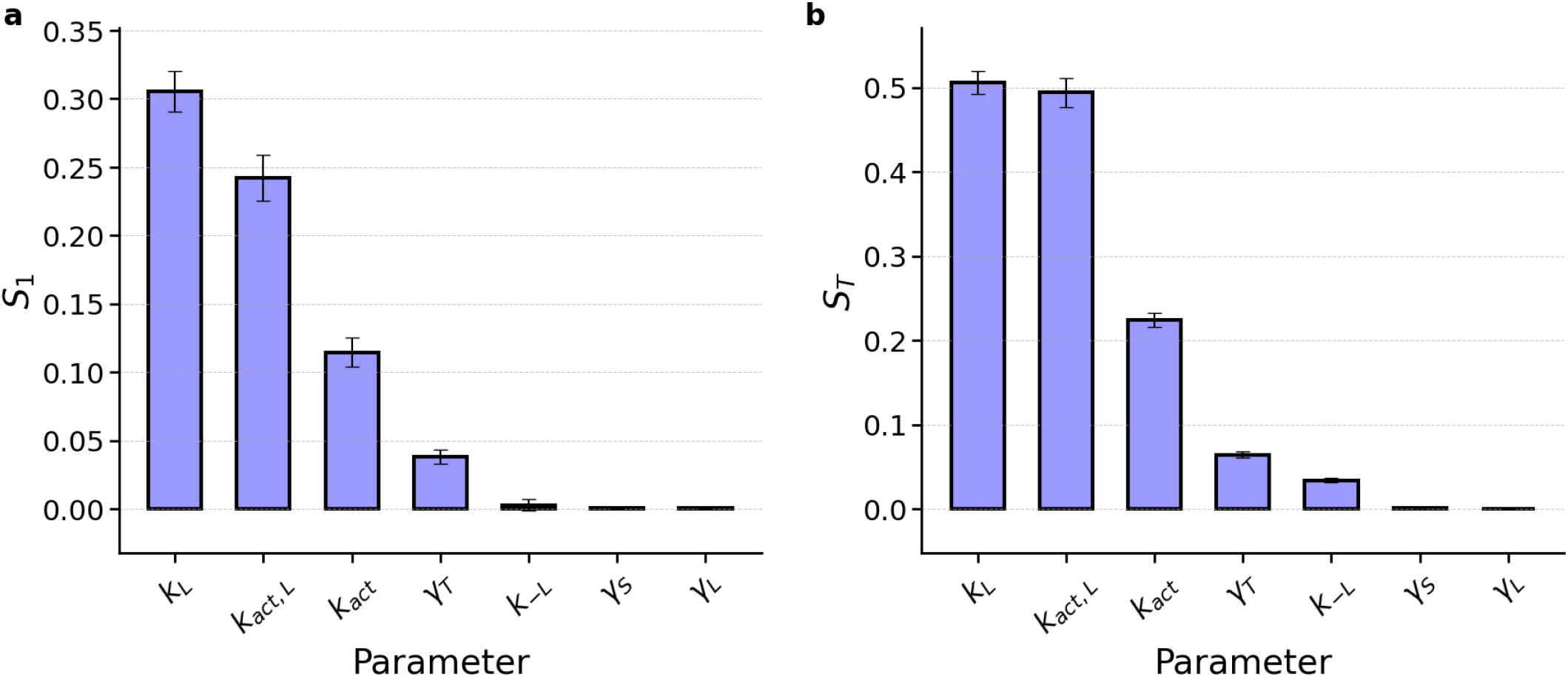
Sobol sensitivity results for the synNotch model. **a**. SynNotch *S*_1_ Sobol results for parameters varied three orders of magnitude from their base values, with 2^14^ samples. **b**. SynNotch *S*_*T*_ Sobol results for parameters varied three orders of magnitude from their base values, with 2^14^ samples.

We also predict that ligand-independent activation (*k*_*act*_) is a moderately-sensitive parameter using sensitivity analysis (see Figs. 3a,b). In experiments, one study altered the transmembrane domain by adding a hydrophobic sequence called RAM7 to synNotch, which reduced ligand-independent syn-Notch activation 14.6-fold (30). This experimental evidence thus partially validates our sensitivity analysis result. In addition, we find that parameters such as the bound and unbound synNotch degradation rates (*γ*_*L*_ and *γ*_*S*_, respectively) have low Sobol indices, meaning that our model predicts that varying them in wide ranges around their baseline values does not contribute much variation in the output. These results suggest that engineering the degradation rates would be less useful than engineering other components of the system.

Different synNotch variants have also been proposed and engineered, with the goal of enabling precise tuning of therapeutic activity while minimizing off-target effects. One approach uses different single-chain variable fragments (scFvs) as extracellular domains, which can bind to specific antigens (75). These scFvs can be engineered to create high-affinity binding domains for a target of interest (13, 75). Since engineering these fragments involves swapping the extracellular domain in synNotch, in our model this changes the relationship between the ligand association and dissociation rates, *k*_*L*_ and *k*_−*L*_. We therefore focus next on using the synNotch model in Eq. (1) to predict dynamic properties of different scFvs. In the following, we determine relevant ligand-association rates (*k*_*L*_) corresponding to different scFvs and consider the impact of these parameters in the syn-Notch model to approximate variant synNotch dynamics.

To parameterize these synNotch variants, we first find the dissociation constant values *K*_*d*_, which are commonly reported for extracellular domains in the literature. *K*_*d*_ is defined in terms of association and dissociation rates using 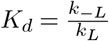.

Since we find that the ligand dissociation rate (*k*_−*L*_) is not sensitive in our model based on the baseline Notch parameters (see Figs. 3a,b), we set this parameter to its constant baseline value as in Table 1 and infer the association rate *k*_*L*_ for each synNotch variant considered. In addition, the Notch-Delta baseline parameters used to generate Fig. 2 using the synNotch model were obtained from a model describing the contact area between two cells, where the spherical shapes of the cells were approximated as two-dimensional circles (56). Since *K*_*d*_ values of scFvs are typically reported in 3-dimensional space in the literature, we need to convert them to 2-dimensional parameters in order to determine the ligand-association rates for our model system. For this, we use the approach proposed in (56) to convert from a 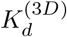 constant to a 2D association rate. First, 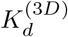 values and 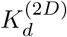 val-ues are related by the characteristic length scale, *λ*_*d*_, as follows (76, 77):

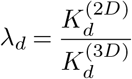

For example, for an anti-CD19 synNotch scFv (78), 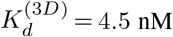. *λ*_*d*_ is assumed to be 1 nm for many biological systems and captures the typical characteristic length scale associated with plasma membrane thickness (76, 77). We thus have the conversion:

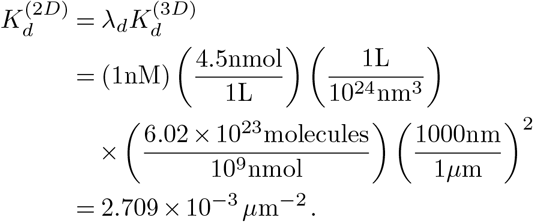

As in (56), we assume that the two-dimensional off-rate 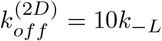 to reflect the case where ligand-dependent activation occurs faster than ligand dissociation, and we therefore obtain 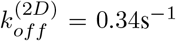. Since 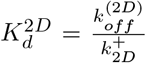, where 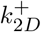 denotes the 2D ligand-association rate, we obtain

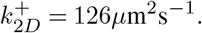

Using this same conversion strategy for 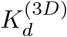 values of different synNotch variants, we then obtain estimated association rates for several specific scFvs, which we report in Table 3.

**Table 3.**
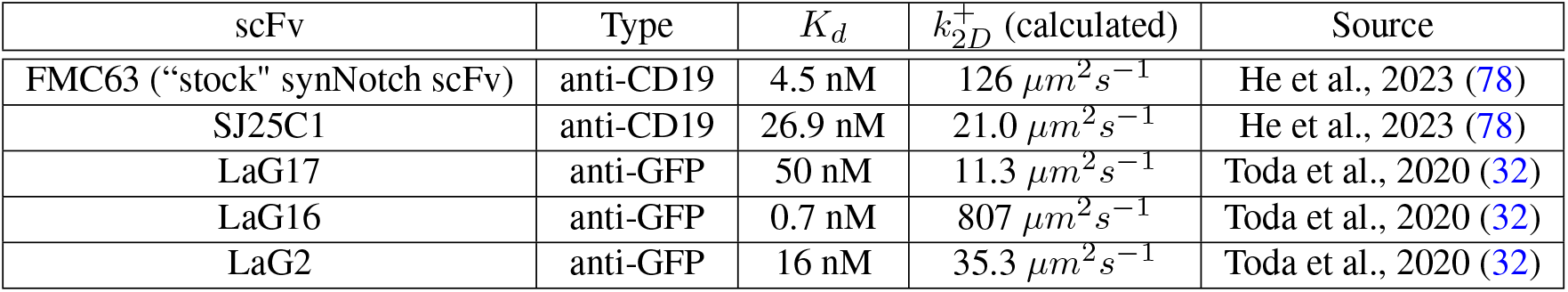
Dissociation constants for various single-chain variable fragments and the corresponding estimated 2-dimensional ligand-association rates.

| scFv | Type | $K_d$ | $k_{2D}^+$ (calculated) | Source |
| --- | --- | --- | --- | --- |
| FMC63 ("stock" synNotch scFv) | anti-CD19 | 4.5 nM | $126 \mu m^2 s^{-1}$ | He et al., 2023 (78) |
| SJ25C1 | anti-CD19 | 26.9 nM | $21.0 \mu m^2 s^{-1}$ | He et al., 2023 (78) |
| LaG17 | anti-GFP | 50 nM | $11.3 \mu m^2 s^{-1}$ | Toda et al., 2020 (32) |
| LaG16 | anti-GFP | 0.7 nM | $807 \mu m^2 s^{-1}$ | Toda et al., 2020 (32) |
| LaG2 | anti-GFP | 16 nM | $35.3 \mu m^2 s^{-1}$ | Toda et al., 2020 (32) |

We use these estimates to simulate the cubic TCM with extracellular variant-specific synNotch parameters. For example, we simulate the anti-GFP LaG17 scFv dynamics by swapping the Notch-Delta association rate (*k*_*L*_) with the calculated one from Table 3 and obtain the predicted dynamics in Fig. 4. The LaG17 dynamics are shown with a dotted pattern, and these results are overlaid onto the Notch-Delta baseline simulations. The results are visualized on a smaller timescale to illustrate the differences in dynamics between the baseline and the variant. The LaG17-synNotch binds much faster to the ligand, as indicated by the larger and steeper *LSynT F* function. Since most synNotch molecules transition to the ligand-bound state quickly, most follow the ligand-dependent activation pathway. This system also saturates much faster than the Notch-Delta system. These differences are relevant on a small timescale, since T-cell interaction times with antigen-presenting dendritic cells have been shown to be as short as one minute before being washed away by blood (79, 80), so these interactions are important on small timescales for predicting binding success.

**Fig. 4.**
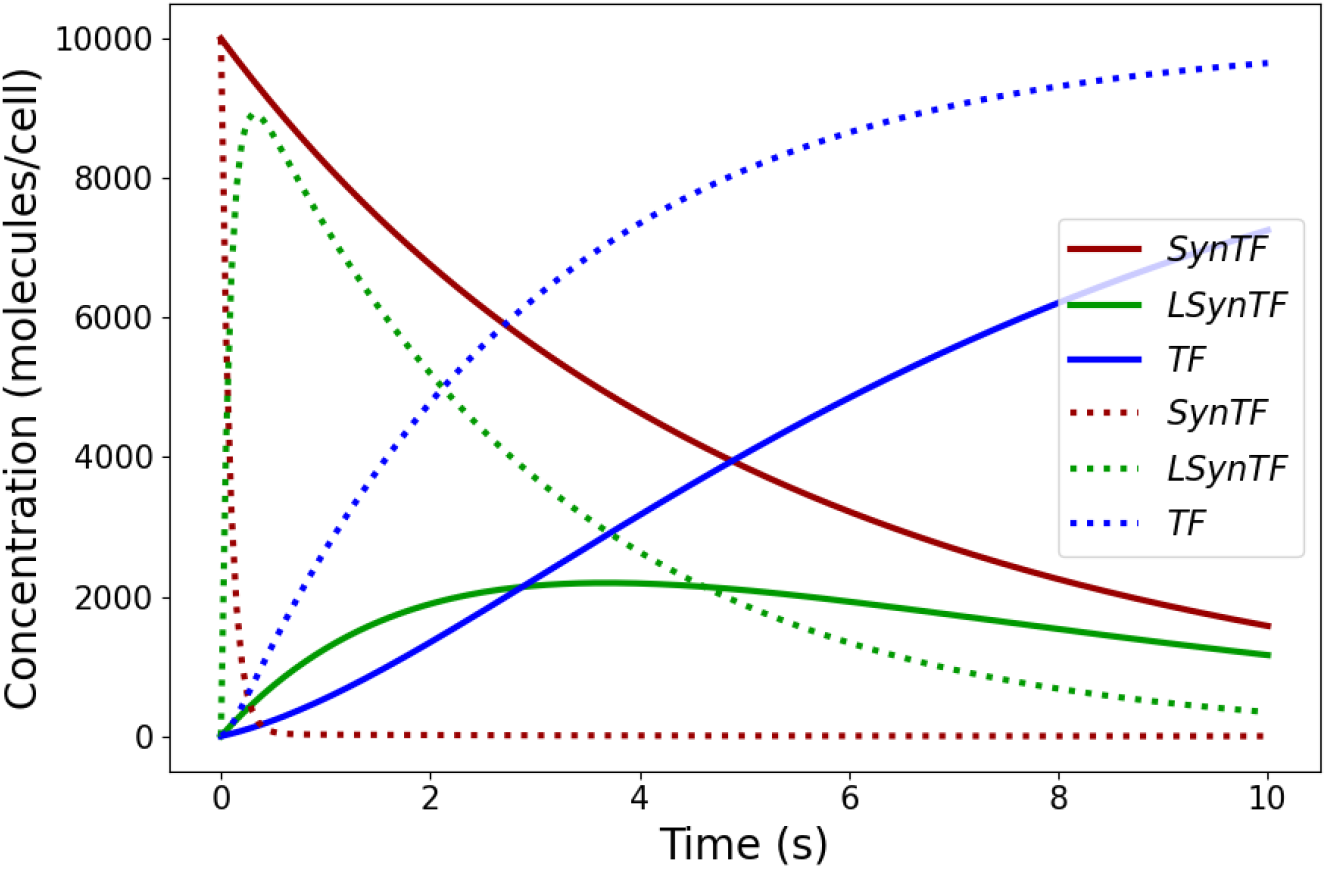
Solution of the cubic TCM with LaG17-synNotch parameters as in Table 3 (dashed lines). Overlaid solution of the cubic TCM with Notch-Delta baseline parameters (Table 1) as in Fig. 2 (solid lines).

### Gene expression model

For the gene expression model in Eq. (2), we first discuss how we estimate several parameter values: the binding energy difference (Δ*ε*), mRNA degradation rate (*γ*_*M*_), mRNA production rate (*r*_*P*_), and fold activation between transcription factor concentration limits (*α*).

We begin by estimating the value of the binding difference Δ*ε*. Previous reports show that the equilibrium expression of GFP under a minimal CMV (minCMV) promoter without induction is roughly 20% as strong as constitutive expression under the CMV promoter, demonstrating that minCMV has non-negligible leaky activation (81). Thus, the probability of RNAP binding to the minCMV promoter without transcription factor induction must be accounted for. To ascertain this binding probability, we first estimate the energy of this interaction. Despite the complexity that underlies RNAP-DNA binding, previous studies have shown that a simple linear model of sequence-dependent binding energy is sufficient to describe RNAP and DNA binding energies (82). In this linear model, each nucleotide in the RNAP binding site contributes a small amount of binding energy, which sums together to approximate the total binding energy of RNAP to that site. Biophysical studies have inferred all 4 *×* 41 parameters for this model, representing every possible nucleotide combination in the −41 to −1 sites (83). Given this data, the binding energy of RNAP to the minCMV promoter can be calculated by adding the appropriate energy values for each nucleotide at each position (83, 84).

The location of the transcriptional start site for this promoter was provided by Isomura et al., 2008 (85). This 4 *×* 41 matrix is reported in *k*_*B*_*T*_*p*_ energy units, and the values are shifted by a constant such that the average binding energy of RNAP across the *E. coli* genome is zero (84). This matrix is available in the Text S2 file of Brewster et al., 2012 (84), and the minCMV promoter sequence is given in the Supplementary material. By adding the values in this matrix corresponding to the promoter nucleotide sequence, we estimate that Δ*ε* = −3.74*k*_*B*_*T*_*p*_, which means that the binding energy of RNAP to minCMV is approximately 3.74*k*_*B*_*T*_*p*_ below the average nonspecific RNAP binding interaction at physiological temperature.

We estimate the half-life of the mRNA transcript using the median mRNA half-life from a large database of mRNAs in human cells, which is 10 hours (62). From this, the mRNA degradation rate is simply given by:

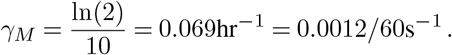

The rate of mRNA production in the state where RNAP is bound (*r*_*P*_) can be calculated using the extension rate of RNAP II in human cells (*r*_*ext*_) and the length of the gene product of interest (*l*):

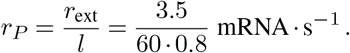

This gives the mRNA production rate in molecules per second. While different CARs have different transcript lengths, we assume a base value in a biologically-relevant range, namely 0.8 kb.

The values of *γ*_*M*_ estimated above, together with *γ*_*C*_ from Table 2, set the timescale for this model on the order of hours. This contrasts sharply with synNotch activation, which occurs on the order of seconds (see Table 1). This allows us to obtain the long-time dynamics of gene expression while approximating the fast dynamics of synNotch. We leverage this fact to estimate two key values in the dynamics of protein production: the equilibrium protein concentration in the absence of induction (*C*_0_), where *T* is absent, and the equilibrium protein concentration at maximal induction (*C*_*h*_), where *T* is maximal. The ratio of these values, together with experimental data, allow for the determination of the unknown parameter *α*. We begin by obtaining the respective mRNA levels *M*_0_ and *M*_*h*_. Based on Eq. (2), the steady-state for mRNA is achieved when:

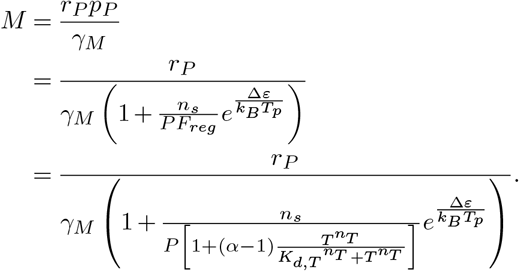

This expression is a function of transcription factor *T*, which changes over time. However, gene expression happens at a much slower timescale than receptor activation, so only steady-state values of *T* are relevant for this analysis. We consider two possible steady states: one where *T* = 0, giving rise to *M*_0_, and one where *T* is very large, giving *M*_*h*_. When *T* = 0, we obtain:

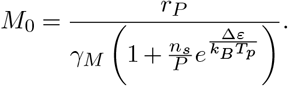

Note that of the parameters specific to engineering synNotch-CAR T-cells, only Δ*ε* and *γ*_*M*_ appear here. Thus, this equilibrium is completely independent of synNotch activation and transcription factor cooperative binding; it is solely a function of the leaky promoter expression and subsequent degradation of product.

To obtain *M*_*h*_, we remove the dependence on *T* of the original *M* equation by instead assuming that *T* equilibrates to a value much larger than *K*_*d*_. This saturates the Hill function dependence on *T* in the *F*_*reg*_ expression in Eq. (2), yielding:

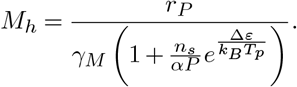

Intuitively, *M*_*h*_ corresponds to the theoretical maximum amount of mRNA that our system can yield. We then calculate the corresponding values of *C*_0_ and *C*_*h*_. For the translated CAR protein in Eq. (2), we reach a steady state when:

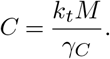

Using the steady states calculated for transcribed mRNA, *M*_0_ and *M*_*h*_, we obtain:

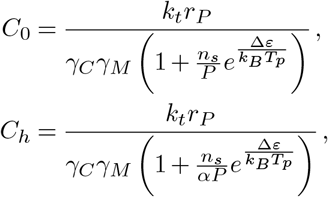

for the cases where *T* = 0, and where *T* is saturating, respectively.

We now take the ratio of the calculated protein levels to compare to a corresponding experimental setting, the details of which are described in the Supplementary material. We find that the steady-state concentration of protein with saturating *T* is 66.97 times larger than the concentration of protein with *T* = 0. Thus, we estimate that:

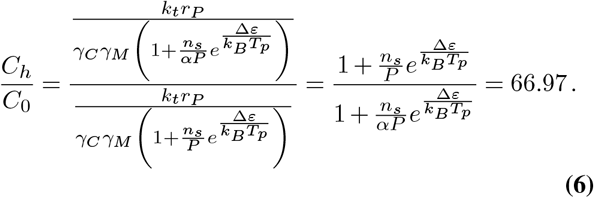

Finally, we can substitute in known values of *n*_*s*_, *P*, and Δ*ε* (see Table 2) to obtain *α* = 110. This ratio is comparable to what has been observed for this promoter in prior quantitative analyses (81).

We now discuss the inclusion of initial synNotch molecules as a gene expression model parameter. The gene expression model is solved on a much longer timescale, so we consider *t* = 0 to reflect when all the transcription factor has been released from synNotch. The degradation rate of the transcription factor is small and will not have a significant effect on the shorter synNotch model timescale, so we adapt the differential equation for *T* (*t*) in Eq. (1) as follows:

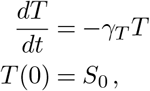

where *S*_0_ is the initial number of synNotch receptors, since each receptor has one transcription factor. This means that 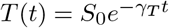, which can be directly substituted into the differential equations for *M* and *C* in Eq. (2) for the gene expression model. This allows us to use the number of syn-Notch receptors as a parameter in the Sobol sensitivity analysis for this model. The results of the dynamic simulation are shown in Fig. 5, where mRNA and CAR concentrations are predicted over time.

**Fig. 5.**
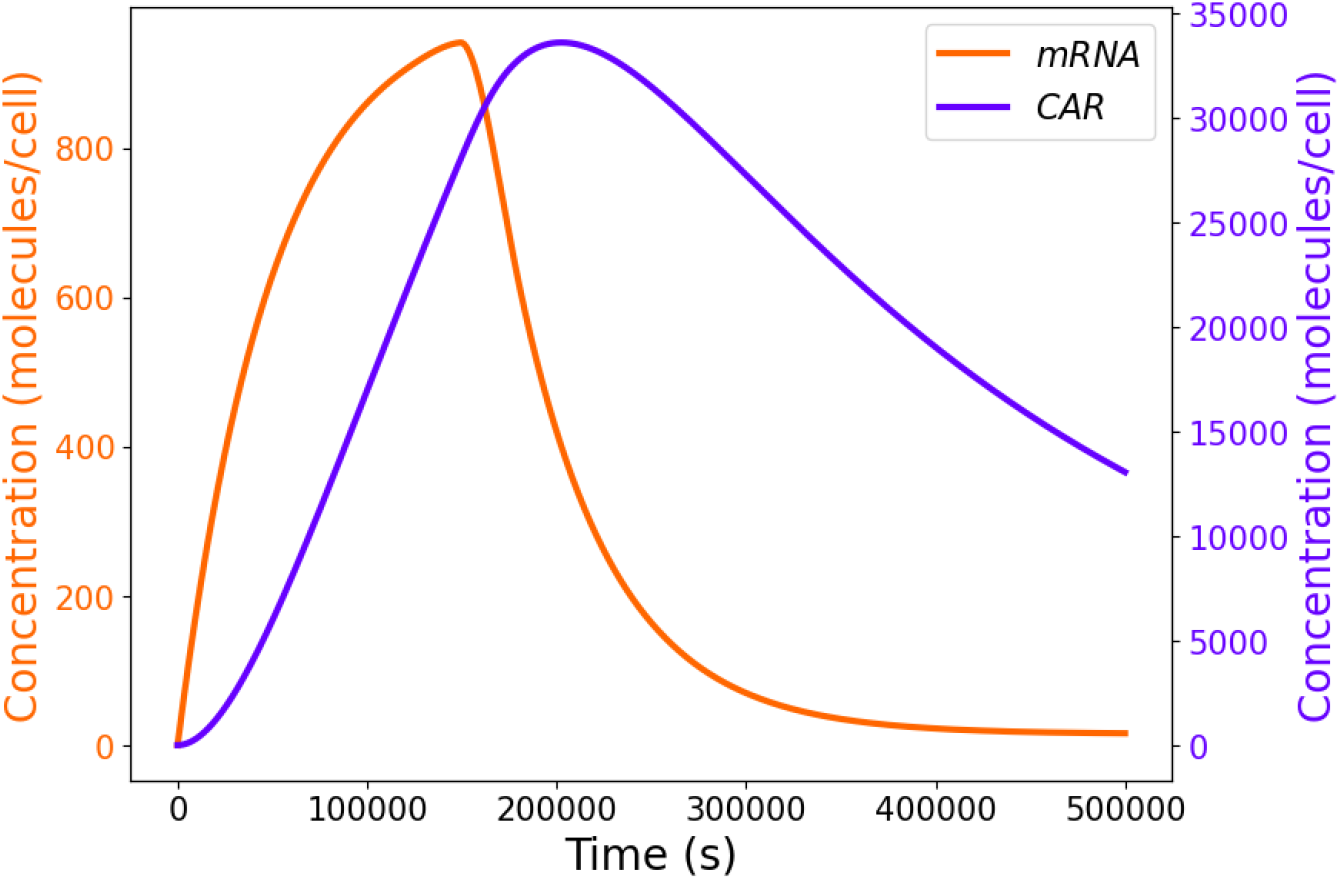
Solution of the gene expression model with gene expression parameters (Table 2).

We then use the *L*^2^-norm of *C*(*t*) used as the output measure for the Sobol sensitivity analysis, on the time interval from 0 to 5 × 10^5^ seconds. The gene expression model parameters are chosen randomly on a logarithmic scale between two orders of magnitude below and two orders of magnitude above their baseline values as shown in Table 2, and we use a total of 2_16_ parameter samples. We only vary the Hill coefficient *n*_*T*_ by one order of magnitude, since biological cooperativity for gene expression and binding events are unlikely to exceed these bounds. The results are shown in Figs. 6a-b. The time period of the simulations allows the solution for *C*(*t*) to increase and then subsequently decrease, as in Fig. 5. While this does not necessarily result in saturation of the concentration of *C*, it is still a useful time frame to compare the impact of different sets of parameter values on the output.

**Fig. 6.**
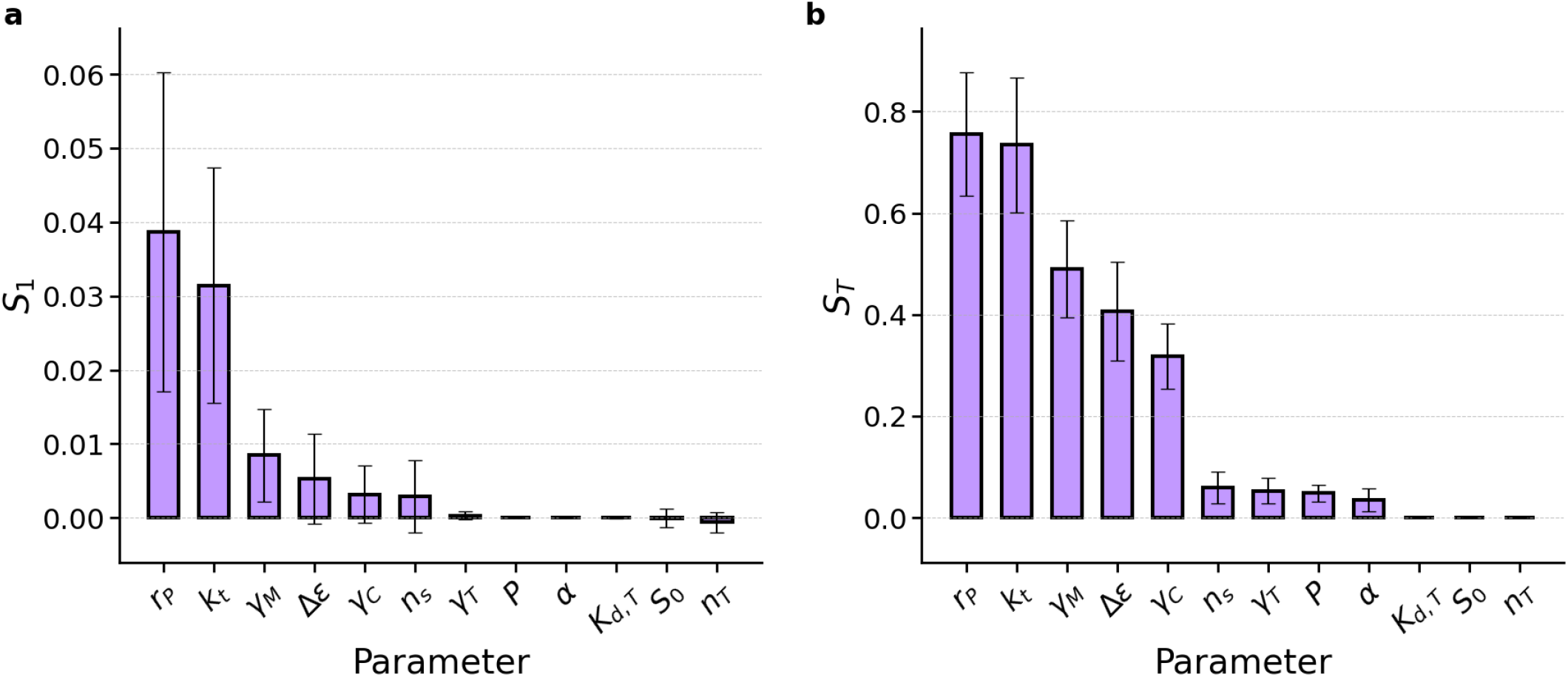
Sobol sensitivity results for the gene expression model. **a**. Gene expression *S*_1_ Sobol results for parameters varied two orders of magnitude from their base values (*n*_*T*_ one order of magnitude), with 2^16^ samples. **b**. Gene expression *S*_*T*_ Sobol results for parameters varied two orders of magnitude from their base values (*n*_*T*_ one order of magnitude), with 2^16^ samples.

We again observe that the computed *S*_*T*_ Sobol indices are larger than the *S*_1_ indices for most parameters. This implies that the interaction terms between parameters have a significant influence on the variability of the output of gene product over time. As a result, we primarily consider the *S*_*T*_ parameters to determine the best engineering targets for dynamic tunability in the gene expression model.

While many synNotch model parameters can be tuned by swapping different component domains, the gene expression model contains some parameters that cannot be directly engineered; that is, certain parameters are more reflective of the cell’s intrinsic machinery and cannot be changed by altering synNotch and CAR alone. The two parameters with the largest *S*_*T*_ indices are the transcription rate with bound RNAP (*r*_*P*_) and the translation rate per mRNA (*k*_*t*_). These two processes are properties of RNAP transcription and ribosome translation in the cell and cannot be readily engineered in a synNotch-CAR T-cell system given the choice of cellular chassis, the host for an engineered biological system.

However, the following three most sensitive parameters shown in Fig. 6b are the mRNA degradation rate (*γ*_*M*_), the RNAP-DNA binding energy difference (Δ*ε*), and the CAR degradation rate (*γ*_*C*_). Engineering degradation rates of mRNAs has been demonstrated by adding AU-rich elements (86). In addition, engineering degradation rates of gene products is possible using various techniques, such as adding degradation sequences like N-degrons and C-degrons (87) and auxin-inducible degrons (88). The binding energy difference can be modified by using a promoter with a different strength, which is a readily available engineering tool (84, 89). The next four parameters listed in Fig. 6b have small positive *S*_*T*_ indices, indicating that they are less sensitive than the other parameters discussed thus far. The degradation rate of the transcription factor (*γ*_*T*_) is not particularly sensitive, indicating that this should not be the first experimental trial when attempting to modulate gene expression dynamics. The number of nonspecific binding sites for RNAP (*n*_*s*_), number of RNAP molecules (*P*), and maximum fold change in available RNAP with saturating transcription factor (*α*) are not readily tunable.

The final three parameters listed in Fig. 6b contribute very little to the overall variation in gene expression dynamics. These final characteristics are important results for engineers. The dissociation constant (*K*_*d*_) and Hill coefficient (*n*_*T*_) for the interaction between the transcription factor and DNA are not sensitive, indicating that attempting to make changes to this interaction may not influence the overall dynamics. Also, the initial number of synNotch receptors does not seem to influence the gene output, so modulating this parameter through promoter engineering is not predicted to yield desired results for maximal activity of synNotch systems. Our mathematical models predict that these parameters with low sensitivity indices will not influence the overall gene expression dynamics as significantly, so they should not be prioritized in engineering experiments.

## 4. Discussion

The complex interactions of synNotch with its extracellular and intracellular environments can lead to undesired responses during the engineering design-build-test-learn cycle, and engineers continue to modify synNotch components to alter its properties for desired effects. Furthermore, the sample space of engineerable options in this system is very large, motivating the use of mathematical modeling to reduce the complexity in the design of experiments (90). Making uninformed changes can incur high experimental costs and yield few promising results in sampling the relevant sequence landscape for synNotch and other engineered receptor systems. Here, we propose mathematical models that can inform engineering decisions, improving on trial-and-error methods that are common in synthetic biology. By understanding which model parameters are the most sensitive and tunable, this work can suggest how to navigate this large parameter space and identify the experiments with the most potential to vary synNotch-driven activation.

We develop a model for synNotch binding and activation and the downstream gene expression it encodes. The synNotch model occurs on a fast timescale, so its kinetics are relevant when considering transient cell-cell interactions. We carry out a global sensitivity analysis of the model, which indicates the parameters that are most likely to yield variation in the synNotch response. We predict that the ligand-association rate between synNotch and its target has the most significant influence on transcription factor release in the system. This also allows us to predict transcription factor dynamics for different dissociation constants *K*_*d*_, which reflect a property of the extracellular domain of synNotch variants. Since different synNotch variants for the same antigen can have different *K*_*d*_ values (32), this is a valuable model output, since it informs engineers about the most appropriate variant for the desired cellular response. In CAR T-cell therapy, the most effective binder is not necessarily the one with the highest affinity (91, 92), underscoring the need for computational tools that can predict optimal binding strength within specific cellular contexts. We also predict that the model is sensitive to ligand-independent activation, which is relevant since engineering efforts have been successful in reducing this activation pathway (30). Since this activation pathway represents receptor leakiness, this is an important factor to control for in immunotherapies like CAR T-cell therapy in order to reduce potentially toxic off-target effects. These parameters can be tuned via experimental and structural adjustments, thus our sensitivity analysis has direct applications to biomedical engineering involving cell and gene therapy optimization. Parameters with lower sensitivity indices also provide valuable insight, as they may suggest experiments which do not yield significant variability in the cellular responses being observed. However, choices such as cellular chassis, copy number variation, and epigenetic silencing of synthetic constructs may alter engineered responses even if not predicted by this model (93–95).

We then couple this model to a general model of gene expression, which predicts the synthesis of a gene product over time. This has applications in CAR T-cell therapy, as syn-Notch activation can be coupled to CAR activation to yield more specific immunological responses. This model occurs on a slower timescale, which is relevant for protein expression. We also carry out global sensitivity analysis of this system, which suggests parameters that could potentially be tuned experimentally to change the gene expression dynamics of the system. Most notably, the promoter strength and the degradation rates of mRNA and CAR are the most sensitive parameters that are also tunable, suggesting that these characteristics should be prioritized when planning experiments to modulate synNotch-CAR gene expression dynamics. With additional experimental validation, this model has powerful predictive capabilities, generating new avenues for engineering in synthetic biology.

Furthermore, our model for synNotch can be directly integrated with existing mathematical models for CAR. The receptor-antigen interaction plays a critical role in CAR models, and antigen binding is explicitly represented through model parameters in existing models (96, 97). Our model can be used when synNotch scFvs mediate these interactions via *k*_*L*_ rates specific to these variants. This multi-receptor synNotch-CAR T-cell system could overcome challenges posed by antigen heterogeneity by using receptor AND/OR-gate logic.

The synNotch-CAR model presented here and its derivation from the cubic TCM could be translated to other systems where a synthetic receptor is tied to gene expression. Other synthetic transmembrane receptors include synthetic intramembrane proteolysis receptors (SNIPRs) and modular extracellular sensor architecture (MESA), which also follow the mechanism of ligand binding, cleavage, and transcription factor release (98–100). These systems use different architectures and are engineered for different functions, but their shared mechanistic properties may allow for the derivation of similar informative mathematical models and sensitivity analyses.

Many models in mathematical biology have parameters that reflect intrinsic properties of the system and may not change. Performing sensitivity analysis in these models can improve understanding of the system, but may not be directly applied to perturb the system. The application we consider here is that of synNotch-activated CAR T-cells, which is a biological system that can be engineered through protein engineering, computation-aided design, and evolution strategies. Here, realistic modeling and rigorous sensitivity analysis provide clear predictions for informing experimental directions and engineering decisions. Our approach provides a rigorous quantitative foundation for modeling and understanding the synNotch-CAR T-cell system, and sensitivity analysis of these models reveals promising avenues for the engineering of synNotch and of the CAR expression response it induces. We hope that this model can be utilized in future engineering cycles for predicting tunable experimental parameters and targets.

## ACKNOWLEDGEMENTS

The authors acknowledge the Bass Connections program at Duke University, the Pratt School of Engineering, the Center for Advanced Genomic Technologies, and the Thomas Lord Foundation of North Carolina for their support.

## Supplementary material

### Overview of Sobol sensitivity analysis

We summarize the background for Sobol sensitivity analysis, as introduced in (101). Given a mathematical model with parameters 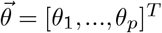 (where *p* is the number of model parameters), we consider an output measure *Y*, which is a function of the parameter vector 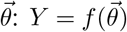. This measure can always be calculated directly from the model given some set of initial parameters. In global sensitivity analysis, each parameter is drawn randomly from a relevant interval. This interval will vary for each parameter, but we can rescale it to the interval [0, 1] for easier analysis and interpretation. To generate a sequence of randomly-distributed points in this unit hypercube of parameters, we use Latin Hypercube Sampling (LHS). This methods splits the parameter space into a grid, and chooses sample parameter values so that there is only one sample in each row and each column of the grid (102).

Sobol decomposition expands 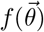 in the following hierarchical way:

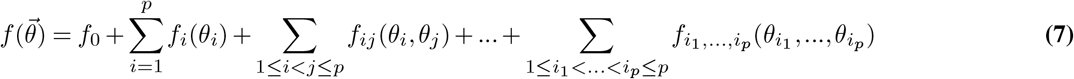

where *f*_0_ is the mean of the function, *f*_*i*_(*θ*_*i*_) are the first-order interactions between parameters, 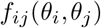 are the second-order interactions between parameters, and 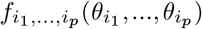 are the *p*^*th*^-order interactions between parameters. This decomposition is not unique, but the interaction term functions can be constrained in the following way:

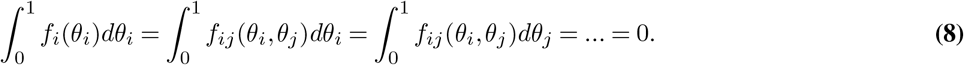

The terms *f*_*i*_(*θ*_*i*_) and the higher-order interaction terms are then computed recursively using Eq. (7) and the constraint in Eq. (8). We can also interpret these integrals in terms of expectations of random variables. In the actual implementation of Sobol sensitivity analysis, the randomly-drawn samples of parameter values are used to assess the variance of the output measure. It can be shown that the first- and higher-order terms have expected values equal to 0. In this probabilistic framework, we can define *D* as the total variance of our measure *Y* :

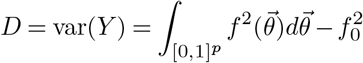

since *f*_0_ = 𝔼 (*Y*). Similar to the decomposition of *f*, the total variance can be decomposed as follows:

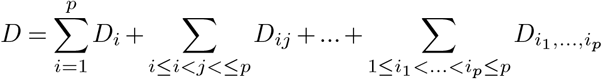

where

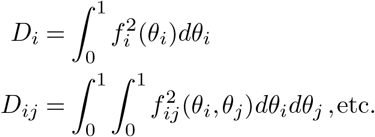

This allows us to define the Sobol indices, which are the main outputs of the Sobol sensitivity analysis. The first-order sensitivity index *S*_*i*_ is defined as

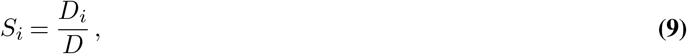

and the total sensitivity index is defined as

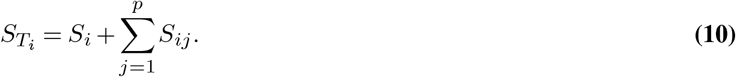

In practice, we only consider up to second-order interactions between parameters. A large *S*_*i*_ value indicates that parameter *i* strongly influences the variance in the measure. A large 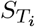 value indicates that parameter *i* and its interactions with other parameters strongly influence the variance in the measure.

### MinCMV promoter sequence

The full minCMV promoter DNA sequence used to calculate the binding energy difference Δ*ε* is shown below.

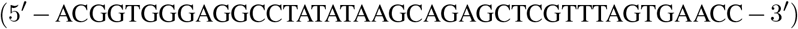

### Experimental protocol

The following experimental protocol from the Kim lab was used to determine the value of the fold-activation for the minCMV promoter, *α*. Two samples of HEK 293T cells were prepared: induced cells and uninduced cells. Induced HEK 293T cells were prepared by co-transfection with pHR_Gal4UAS_tBFP_PGK_mCherry and pcDNA5_Gal4VP64 using Thermo Fisher Lipofectamine. Un-induced HEK 293T cells were prepared by transfection with pHR_Gal4UAS_tBFP_PGK_mCherry using Thermo Fisher Lipofectamine. For all cells, BFP and mCherry fluorescence was quantified by flow cytometry 48 hours post-transfection. The relative activation of induced and un-induced cells is given by median cellular BFP intensity divided by median cellular mCherry intensity, which normalizes for transfection efficiency. The fold-activation is then calculated using the ratio of induced activation to un-induced activation as in Eq. (6).

